# Two time scales of adaptation in human learning rates

**DOI:** 10.1101/2025.06.05.658048

**Authors:** Jonas Simoens, Senne Braem, Pieter Verbeke, Haopeng Chen, Stefania Mattioni, Mengqiao Chai, Nicolas W. Schuck, Tom Verguts

## Abstract

Different situations may require radically different information updating speeds (i.e., learning rates). Some demand fast learning rates, while others benefit from using slower ones. To adjust learning rates, decision makers could rely on either global, meta-learned differences between environments, or faster but transient adaptations to locally experienced prediction errors. Here, we introduce a new paradigm that allows researchers to measure and empirically disentangle both forms of adaptations. Participants performed short blocks of trials of a continuous estimation task – fishing for crabs – on six different islands that required different optimal (initial) learning rates. Across two experiments, participants showed fast adaptations in learning rate within a block. Critically, participants also learned global environment-specific learning rates over the time course of the experiment, as evidenced by computational modelling and by the learning rates calculated on the very first trial when revisiting an environment (i.e., unconfounded by transient adaptations). Using representational similarity analyses of fMRI data, we found that differences in voxel pattern responses in the central orbitofrontal cortex correlated with differences in these global environment-specific learning rates. Our findings show that humans adapt learning rates at both slow and fast time scales, and that the central orbitofrontal cortex may support meta-learning by representing environment-specific task-relevant features such as learning rates.

## Introduction

Decisions that humans make on a daily basis range from the ordinary, such as what to have for lunch, to the life-defining, such as what career to pursue. The computational framework of reinforcement learning (RL) stipulates that such decision making requires estimating and continuously updating relevant feature values of specific options (e.g., the nutritional value, the tastiness, and the price of the lunch), and subsequently using these estimates to make a good choice. The most common RL approach to learning such values is the delta rule (Rescorla & Wagner, 1972):

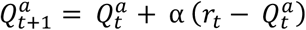

where the (*Q*) value of action *a* at time *t* + 1 is updated in proportion to the error made in predicting the outcome *r* of action *a* at time *t* (i.e., the prediction error). This learning algorithm requires choosing a learning rate *α*. Performance of the algorithm depends on the value of this parameter, and different values are optimal for different environments (Sutton & Barto, 2018; Verbeke & Verguts, 2024). Learning the parameters that shape learning is termed meta-learning (Binz et al., 2024; Wang et al., 2018), and theories of human meta-learning suggest that people can flexibly adapt their learning rates to the environment (Mathys, 2011; Schweighofer & Doya, 2003; Silvetti et al., 2018). Similarly, in artificial agents, setting a learning rate is critical, and a breakthrough in AI was the development of an algorithm that set its learning rate in an adaptive manner (Adam optimizer (Kingma & Ba, 2017)).

Consistent with these ideas, humans can adjust their learning rate to the reward volatility and variability of a given environment (Behrens et al., 2007; Browning et al., 2015; Cook et al., 2019; Goris et al., 2021). An important limiting aspect of these studies, however, is that participants typically spend extended periods of time within one environment. Therefore, previous studies cannot dissociate between fast, transient adaptations (i.e., fast time scale learning within an environment; (Bai et al., 2014; Krugel et al., 2009; Nassar et al., 2012) versus learned environment-specific learning rates that can be reused when revisiting an environment (i.e., slow time scale learning about an environment (Simoens et al., 2024)). From an optimality perspective (Kalman filter; (Dayan et al., 2000)), the learning rate should gradually decrease as the task statistics within an environment become increasingly known. When the environmental statistics are reset (e.g., when visiting a new environment), learning rate should be reset as well. Nevertheless, if the higher-order statistics in an environment remain fixed (e.g., amount of noise in the environment), an optimal agent could (meta-)learn higher-order parameters (e.g., the starting point of the learning rate) so that learning in the environment becomes increasingly efficient. From this perspective, people could learn about the optimal learning rate at two levels: On a fast time scale in response to local prediction errors as an environment becomes known, and on a slower time scale as an environment’s higher-order statistics and optimal initial settings are learned.

To test this, we administered a novel task in two experiments in which participants went fishing for crabs on six different locations around an island that allowed us to measure both trial-by-trial (transient) adaptations in learning rate as a function of locally experienced prediction errors, as well as learned variations in learning rate adapted to the environmental statistics. This task required continuous responses (i.e., estimating the crab locations) and provided continuous feedback (on those crab locations), which allowed us to estimate learning rates on a trial-by-trial basis by calculating how much people updated their fishing location based on feedback (Nassar et al., 2012). Importantly, each of the six locations around the island had one of three different optimal initial learning rates. This was achieved by changing two quantities that governed the outcome (crab location) distributions of each island: the standard deviation of the distribution that determined the latent mean of crab locations upon an island visit (prior distribution), and the standard deviation of the noise with which crabs were sampled around their latent mean (sampling distribution). As a result, different locations around the island required different learning rates for optimal task performance.

Participants switched between locations on a block-by-block basis, where blocks only lasted two to ten trials (depending on the experiment, see below). Crucially, without prior experience, environments were indistinguishable up to the moment feedback was provided on the second trial of each block, so any differences in learning rates between locations after feedback on the first trial suggest meta-learning of environment-specific learning rates. Across two experiments, we found that participants dynamically updated their learning rate within environments, but also, critically, learned over time to use different initial learning rates when revisiting different environments, showing the meta-learning of learning rates tailored to the different environments.

In the second experiment, we also collected fMRI data to investigate which brain regions are involved in representing sustained environment-specific learning rates. Namely, we performed representational similarity analyses on neural voxel pattern activity when participants had just been transported to the next location around the island, while they were preparing to perform the task in a particular environment. We hypothesized that the orbitofrontal cortex (OFC) would be the main region involved in this preparation, consistent with its broader role in the representation of task states (Moneta et al., 2024; Schuck et al., 2016; Stalnaker et al., 2015; Wilson et al., 2014)– the integration of contextual information that is necessary to predict the outcomes of decisions, crucial for reward maximization. While shifts in task state representations and value computations are often linked to the OFC, the environment-specific responses to reward predictions are often linked to the basal ganglia. Therefore, we also studied ventral striatum activity as a candidate for processing prediction errors on the first trial of each block. That is, given the role of the ventral striatum in processing (reward) prediction errors (Calderon et al., 2021; O’Doherty et al., 2004; Pessiglione et al., 2006; Schultz et al., 1997), we assumed that this region would show the effect of these learned environment-specific learning rates through a differential response to on-task (reward) prediction errors.

## Results

### Experiment 1

Fifty participants performed a novel crab-fishing task. At the start of each of 60 blocks, a boat took them to one of six locations around an island (Figure 1A). In each location, participants could drop a cage on a chosen position ten times (trials), with the goal to catch as many crabs as possible (Figure 1C). Each time a cage was dropped, five crabs appeared and spread out evenly from one position in the sand sampled from the sampling distribution 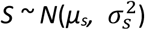, a truncated normal distribution ranging between *µ*_*s*_ *± 1*.*65 * σ*_*s*_. Each crab was either caught by the cage or ran away. At the beginning of each block, the latent mean of the sampling distribution, *µ*_*s*_, was sampled from the prior distribution 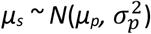, truncated between *µ*_*p*_ *± 1*.*65 * σ*_*p*_, with *µ*_*p*_ set to be the centre of the screen, where the cage position was initialized on the 1st trial of each block. Crucially, the variance of the latent mean 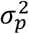, and the noise variance around that mean 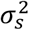, were dependent on the location around the island. In two randomly selected adjacent locations, 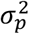 was large, while 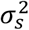 was small. Here, the true mean position of the crabs was widely dispersed and could be nearly everywhere on the screen, while the individual crabs appeared very close to that true mean. Hence, participants could infer the mean crab position from a single observation and performed best if they strongly adjusted the position of the cage after the first trial. We termed this the *low noise environment*, which required a high initial learning rate (Figure 1B). On the two adjacent locations on the exact opposite side of the island, the situation was reversed (i.e., small 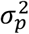, large 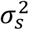, henceforth the *high noise environment*), requiring a low initial learning rate. Finally, on the two locations in between, 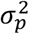 and 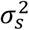, were intermediate and equal (i.e., the *medium noise environment*), requiring an intermediate initial learning rate.

**Figure 1.**
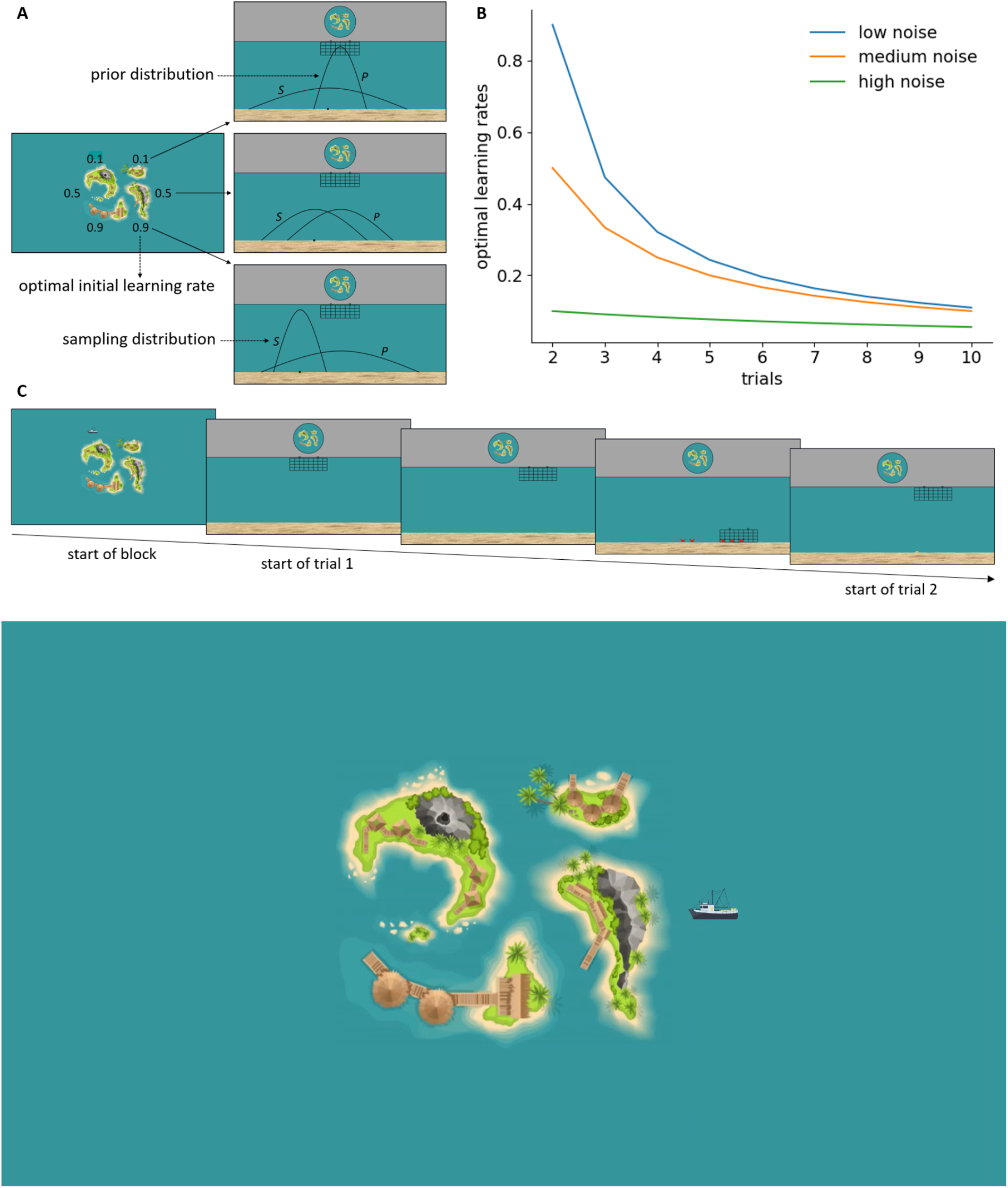
Experimental design. **A:** Participants went fishing for crabs on six different locations around an island which differed in terms of optimal initial learning rate. At the beginning of each block, *µ*_*s*_ was sampled from the prior distribution 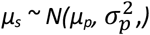, truncated between *µ*_*p*_ *± 1*.*65 * σ*_*p*_, with *µ*_*p*_ = the centre of the screen. Subsequently, on each trial, once a cage was dropped, five crabs appeared and spread out from one location in the sand sampled from the sampling distribution 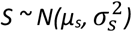), truncated between *µ*_*s*_ *± 1*.*65 * σ*_*s*_, each of which was either caught by the cage or ran away. **B:** Overview of optimal learning rates for performing the task for 1 block of trials according to the Kalman filter assuming that measurement 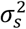 and estimate 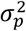 on trial 1 (See *Model estimation and selection* for details). **C:** Overview of the trial procedure (see also Video 1). At the beginning of each block, participants were taken to one of six locations around the island. On each trial, participants positioned the cage somewhere along the x-axis of the screen and dropped it. As the cage sank, five crabs appeared out of one point in the sand and spread out. When the cage reached the ocean floor, crabs caught by the cage remained there while the other crabs ran away. At the start of the next trial, the cage was again at the top of the screen, but at the same x-coordinate where it was dropped in the last trial, and a little heap of sand was left where the five crabs had appeared out of the sand on the last trial.

On the first trial of each block, participants could only drop the cage in the centre of the screen. Crucially, the position of the first crabs was determined by the randomly drawn latent mean (with variance 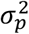), and the variance of the sampling distribution 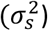, i.e. corresponding to a normal distribution with mean equal to the centre of the screen and variance equal to 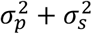. Although 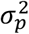 and 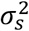 differed across the different environments as described above, their sum was constant, and hence the normal distribution for the first trial in a block was identical across the three environments. Therefore, the first prediction errors participants experienced across the three environments were identical, which we also confirmed by analysing the feedback data. This implies that variations in the first learning rate are uniquely attributable to the meta-learned learning rates.

#### Behavioural results

We first evaluated whether our design was successful in inducing variations in learning rate within environments – where learning rate was defined as the percentage of the direction taken from the last cage drop to last appearance of crabs (see Methods). Specifically, consistent with the optimality analyses depicted in Figure 1B, we assumed that learning rate would decrease over the course of a block, as participants learned more and more about the crab locations within that block. Consistently, a linear mixed effects model analysis, with a random intercept for participant and random slopes for environment and trial number, indicated that learning rates significantly decreased over trials (within blocks) (Figure 2A; *ß* = -0.056, SE = 0.006, *p* < .001).

**Figure 2.**
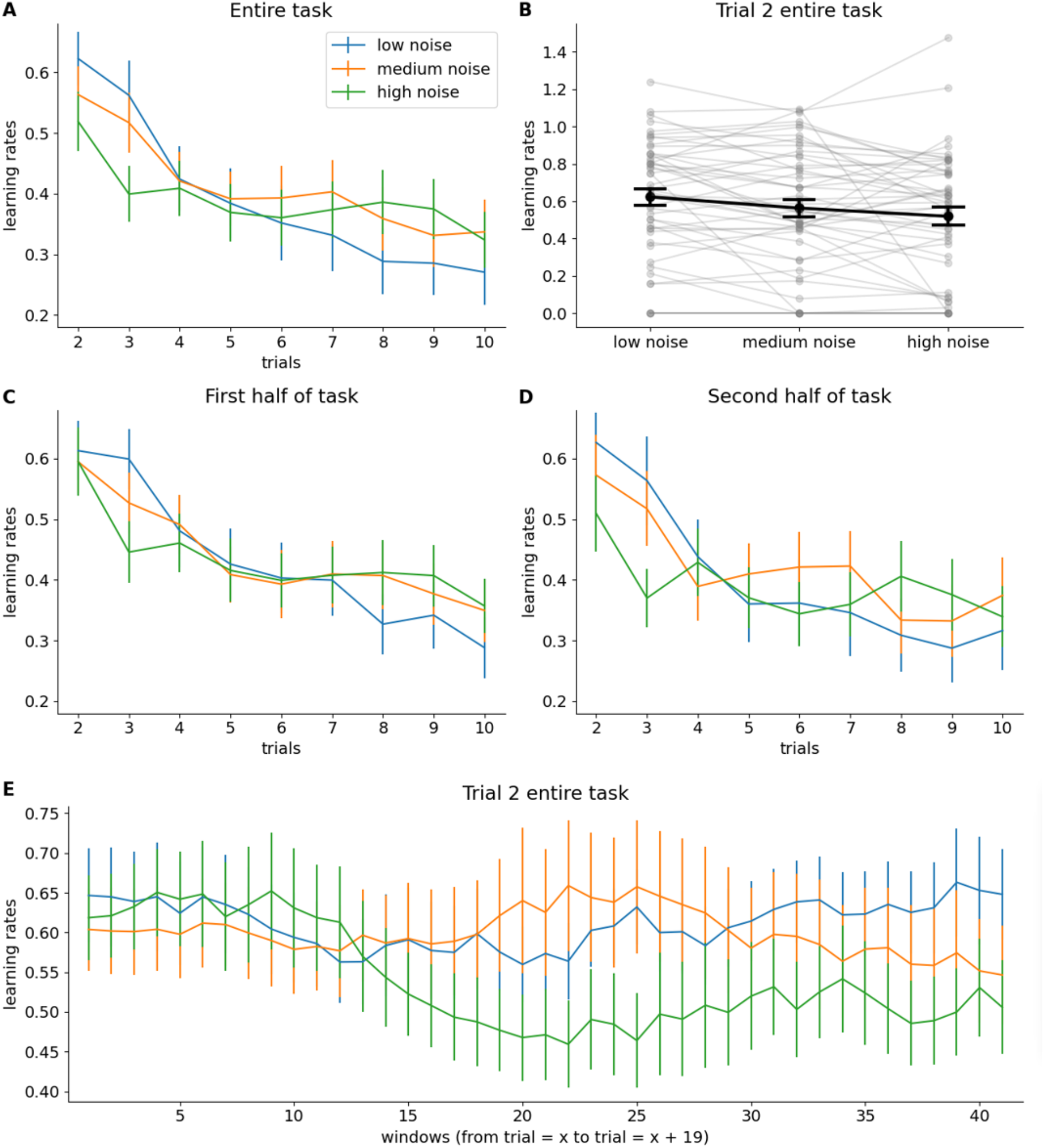
Behavioural results Experiment 1. **A:** Group-level mean of each participant’s median learning rate for each trial in each environment (See *Behavioural data analyses* for details). Error bars represent standard errors of the means. **B**: Detailed overview of all participants’ median initial learning rates. **C-D:** Evolution of (group-level mean) learning rates over trials (within blocks) for the first half (C) and the second half (D) of the task separately. E: Moving-window analysis of trial-2 learning rate across blocks.

Second, to probe whether people were also able to learn and adapt their learning rate to the environment-specific differences, we analysed the learning rates separately focusing on the second trial only (after their first feedback). Differences in learning rates between environments after first feedback could not be attributed to differences in experienced prediction errors, as prediction errors after the first trial were similar across the three environments. Instead, differences in this initial learning rate had to reflect a reaction to learned statistics of the environment from earlier environment visits (i.e., meta-learning). Indeed, we found that observed learning rates on the second trial of each block were significantly higher in the low noise compared to in the high noise environment, *t*(49) = 2.896, *p* = .003 (see Figure 2B). Moreover, these learning rates were also higher in the low noise compared to in the medium noise environment, *t*(49) = 2.407, *p* = .01, but not in the medium noise compared to in the high noise environment, *t*(49) = 1.304, *p* = .099.

Assuming these differences could only have developed over time, after sufficient experience with each of the environments, we also evaluated whether they were more pronounced in the second half of the experiment, compared to the first half. Figure 2C-D indeed suggests this difference in learning rates between the low and high noise environment after the second trial could not yet be observed in the first half of the experiment (*t*(49) = 0.415, *p* = .34), while it appeared in the second half (*t*(49) = 2.067, *p* = .022). However, this was not further corroborated by an interaction between time (first vs. second half of the task) and environment (low vs. medium vs. high noise environment) (*F*(2, 98) = 0.923, *p* = .401). Finally, a moving-window analysis of the 2^nd^-trial learning rate across blocks (with a window size of 20 blocks) did suggest that learning rates gradually decrease in the high-noise condition (see Figure 2E).

#### Modelling results

Next, we turned to computational modelling to evaluate which model could best describe the variations in learning rate reported above. We fitted six models to the data using hierarchical Bayesian analysis (Ahn et al., 2017). We fitted three model types, one of which assumed no local adaptations to learning rates, and two of which allowed for learning rates to be adjusted on a trial-by-trial basis. For each of these model types, we tested both a variant with (initial) learning rates for each environment separately (i.e., environment-specific), and one with a single initial learning rate (non-environment-specific). The first two models were the environment-specific and non-environment specific versions of the Rescorla-Wagner model. The Rescorla-Wagner model assumes that participants updated their *µ*_*s*_ estimates (within blocks) using the delta rule with a fixed learning rate (across trials, within blocks). The next two models were the environment-specific and the non-environment-specific version of the Kalman filter. The Kalman filter assumes that participants updated their *µ*_*s*_ estimates (within blocks) using the delta rule, where the learning rate is a function of estimation noise and measurement noise. Because estimation noise gradually decreases in a block (as people are increasingly aware of the mean crab location), learning rate gradually decreases. Here, initial estimation noise is the free, estimated parameter. The final two models were the environment-specific and non-environment-specific versions of a model that allowed for local, prediction-error weighted changes to the learning rate, here referred to as the Bai model (Bai et al., 2014; Simoens et al., 2024). The Bai model also assumes that participants updated their *µ*_*s*_ estimates (within blocks) using the delta rule. The Bai model further assumes that participants start with an initial learning rate that they up- and down-regulate (within blocks) in proportion to experienced prediction errors. Here, both initial learning rate and decay rate are free, estimated parameters, and both were either environment-specific or non-environment-specific.

According to the leave-one-out information criterion (LOOIC) (Vehtari et al., 2017), the environment-specific Bai model fitted the data best (Table 1), indicating that participants indeed learned to use environment-specific initial learning rates and that they indeed decreased their learning rates over trials (in contrast to what the RW model predicts), but that they did so driven by experienced prediction errors rather than in a statistically optimal way (in contrast to what the Kalman filter predicts).

**Table 1.** Model comparison.

| Model | LOOIC | SE | ΔLOOIC | ΔSE |
| --- | --- | --- | --- | --- |
| Environment-specific Bai model | 28735 | 404 | 0 | 0 |
| Non-environment-specific Bai model | 28582 | 441 | 153 | 65 |
| Environment-specific Rescorla-Wagner model | 28436 | 398 | 299 | 67 |
| Non-environment-specific Rescorla-Wagner model | 28172 | 440 | 563 | 112 |
| Environment-specific Kalman filter | 27853 | 493 | 882 | 211 |
| Non-environment-specific Kalman filter | 27698 | 521 | 1037 | 211 |

*Note.* Models are ranked in descending order according to how well they fit the data. LOOIC refers to a model's approximated expected log pointwise predictive density. Higher values indicate higher out-of-sample predictive fit. SE refers to the standard error of a model's LOOIC. $\Delta$ LOOIC refers to the difference between a model's LOOIC and the top ranked model's LOOIC. $\Delta$ SE refers to the standard error of the difference between a model's LOOIC and the top ranked model's LOOIC.

The posterior probabilities that the initial learning rates (Figure 3A; estimated by the environment-specific Bai model) were higher in the low noise compared to the high noise environment was 0.999; in the low noise compared to the medium noise environment was 0.962; and in the medium noise compared to the high noise environment was 0.999. Decay rates were not significantly different from each other (Figure 3B).

**Figure 3.**
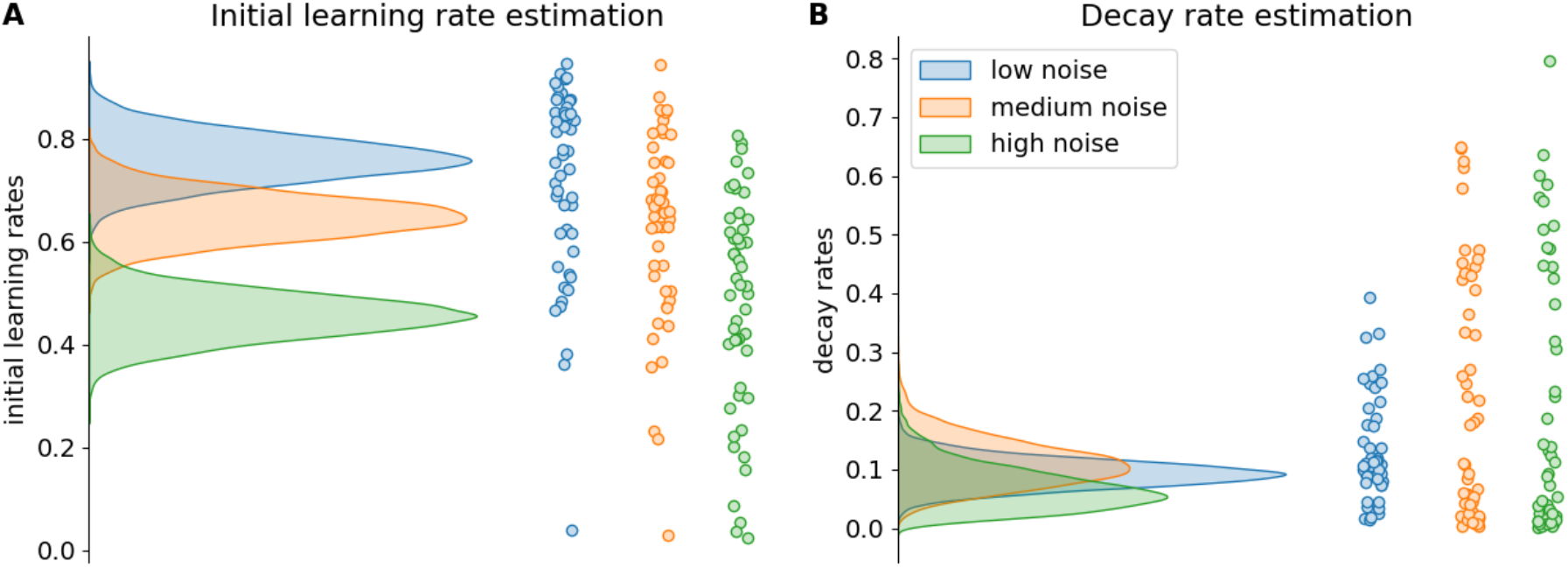
Bai model estimation results Experiment 1. The density plots on the left side of each subfigure show the full posterior densities over the means of the group-level distributions of the relevant parameters. The scatter plots on the right side of each subfigure show the means of all individual-level posterior distributions of the relevant parameters.

### Experiment 2

To establish whether our findings could be reproduced in an independent sample and to investigate where in the brain environment-specific learning rates are represented, we ran a near-exact replication of our first experiment, which 53 participants performed inside an MR-scanner. Experiment 2 consisted of 60 blocks that were identical to the blocks in Experiment 1, except that they consisted of only eight trials. Additionally, 60 blocks consisting of only two trials were randomly intermixed with the 60 longer blocks. These shorter blocks were included to increase power for analysis on the first few trials within blocks. Furthermore, we equipped the fishing boat with a laser pointer (i.e., a vertical red line from the middle of the cage to the sand) so participants could estimate more precisely where their cage would land when dropped. Similarly, to avoid miscalibrations in relation to their previous attempt, we reminded participants of their last cage location by placing a red cross wherever their cage last appeared.

#### Behavioural results

As in Experiment 1, we found that learning rates significantly decreased over trials within blocks (Figure 4A; *ß* = -0.059, SE = 0.008, *p* < .001). Also, observed learning rates on the second trial of each block were significantly different across environments (Figure 4B; *F*(2, 104) = 12.839, *p* < .001). That is, they were significantly higher in the low noise than in the high noise environment (*t*(52) = 3.843, *p* < .001), in the low noise than in the medium noise environment (*t*(52) = 1.998, *p* = .025), and in the medium noise than in the high noise environment (*t*(52) = 3.555, *p* < .001).

**Figure 4.**
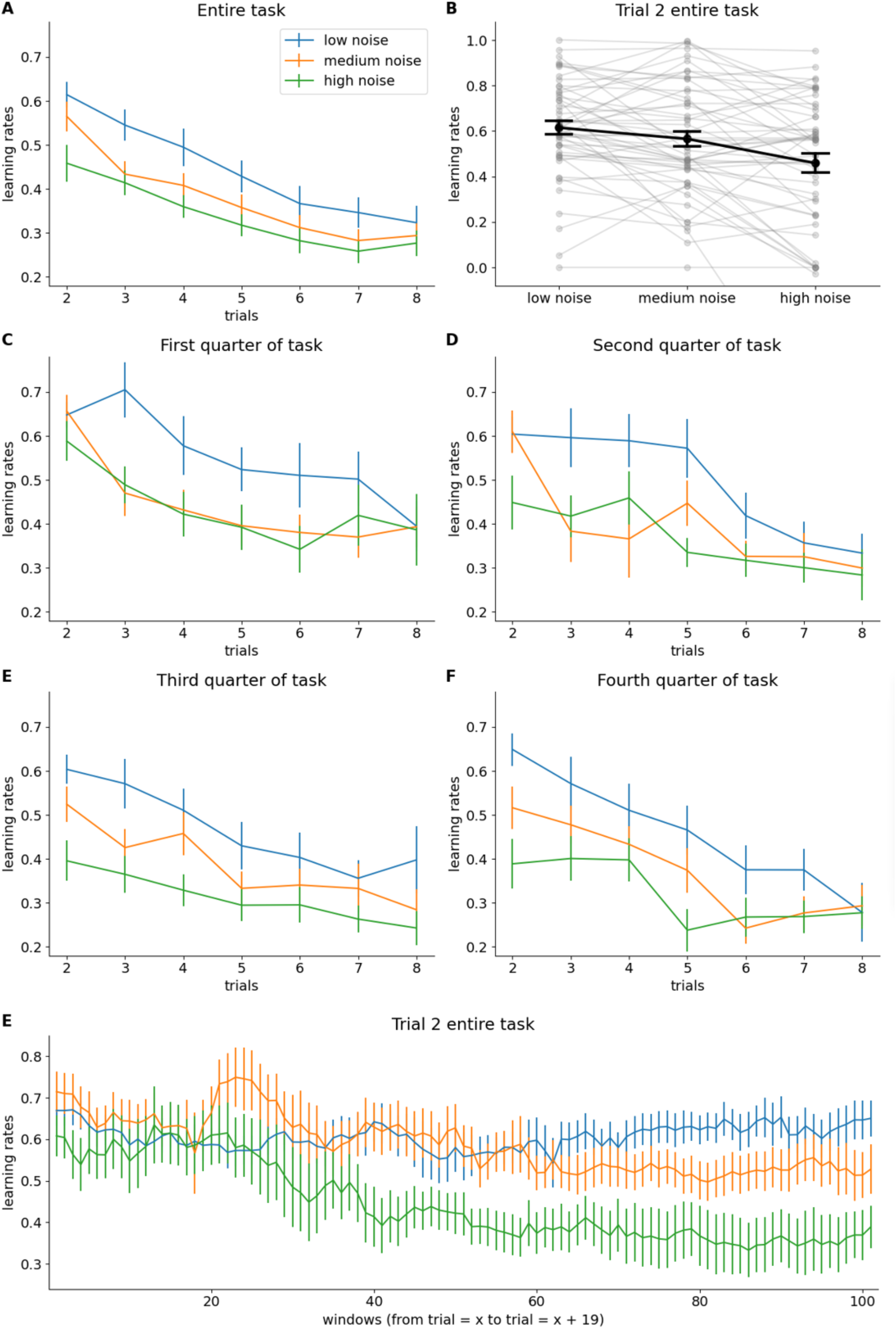
Behavioural results Experiment 2. **A:** Group-level mean of each participant’s median learning rate for each trial in each environment (See *Behavioural data analyses* for details). Error bars represent standard errors of the means. **B**: Detailed overview of all participants’ median initial learning rates. One participant’s median initial learning rate of -0.674 in the high noise environment is not visible on the plot. **C-F:** Evolution of (group-level mean) learning rates over trials (within blocks) for each quarter of the task separately. **G**: Moving-window analysis of 2^nd^-trial learning rate.

Because we had twice as many first trials as in Experiment 1, we next compared quarters of the task rather than halves of the task as done in Experiment 1, as a more sensitive measure of what they learned over the course of the experiment. In line with our expectation that initial learning rates reflect learned statistics of the environment, a significant interaction between time (four quarters of the tasks) and environment (three measurement noises) indicated that learning rates on the second trial of each block became more environment-specific over time (Figure 4C-E; *F*(6, 312) = 2.819, *p* = .011). To unpack this interaction, in the first quarter, initial learning rates were significantly higher in the medium noise environment than in the high noise environment (*t*(52) = 2.252, *p* = .014), but not in the low noise compared to the high noise environment (*t*(52) = 1.338, *p* = .093), nor in the low noise compared to in the medium noise environment (*t*(52) = -0.258, *p* = .601). In the second quarter, initial learning rates were significantly higher in the low noise than in the high noise environment (*t*(52) = 2.442, *p* = .009) and in medium noise compared to in the high noise environment (*t*(52) = 2.443, *p* = .009), but not in the low noise compared to in the medium noise environment (*t*(52) = - 0.12, *p* = .548). In the third and fourth quarters, initial learning rates were significantly higher in the low noise compared to in the high noise environment (*t*(52) = 4.164, *p* < .001; *t*(52) = 4.116, *p* < .001), in the low noise compared to in the medium noise environment (*t*(52) = 2.207, *p* = .016; *t*(52) = 2.829, *p* = .003), and in the medium noise compared to in the high noise environment (*t*(52) = 3.073, *p* = .002; *t*(52) = 2.425, *p* = .009). Finally, the same moving-window analysis again suggests a gradual decrease of learning rate in the high-noise condition, and smaller decrease in the medium-noise condition (Figure 4G).

#### Modelling results

Replicating our findings from Experiment 1, the environment-specific Bai model fitted the data best (Table 2), indicating that participants indeed learned to use environment-specific initial learning rates and that they indeed decreased their learning rates over trials (as opposed to what the RW model predicts), but that they did so driven by experienced prediction errors rather than in a statistically optimal way (as opposed to what the Kalman filter predicts).

**Table 2.** Model comparison.

*Model comparison*
| Model | LOOIC | SE | ΔLOOIC | ΔSE |
| --- | --- | --- | --- | --- |
| Environment-specific Bai model | 34725 | 142 | 0 | 0 |
| Non-environment-specific Bai model | 34682 | 158 | 43 | 34 |
| Environment-specific Kalman filter | 34592 | 160 | 133 | 42 |
| Environment-specific Rescorla-Wagner model | 34567 | 136 | 158 | 32 |
| Non-environment-specific Kalman filter | 34523 | 174 | 202 | 49 |
| Non-environment-specific Rescorla-Wagner model | 34434 | 154 | 291 | 51 |

*Note.* Models are ranked in descending order according to how well they fit the data. LOOIC refers to a model's approximated expected log pointwise predictive density. Higher values indicate higher out-of-sample predictive fit. SE refers to the standard error of a model's LOOIC. $\Delta$ LOOIC refers to the difference between a model's LOOIC and the top ranked model's LOOIC. $\Delta$ SE refers to the standard error of the difference between a model's LOOIC and the top ranked model's LOOIC.

The posterior probabilities that initial learning rates estimated by the environment-specific Bai model (Figure 5A) were higher in the low noise compared to the high noise environment, in the low noise compared to the medium noise environment, and in the medium noise compared to the high noise environment, were 0.999, 0.992, and 0.992, respectively. Decay rates were not significantly different (Figure 5B).

**Figure 5.**
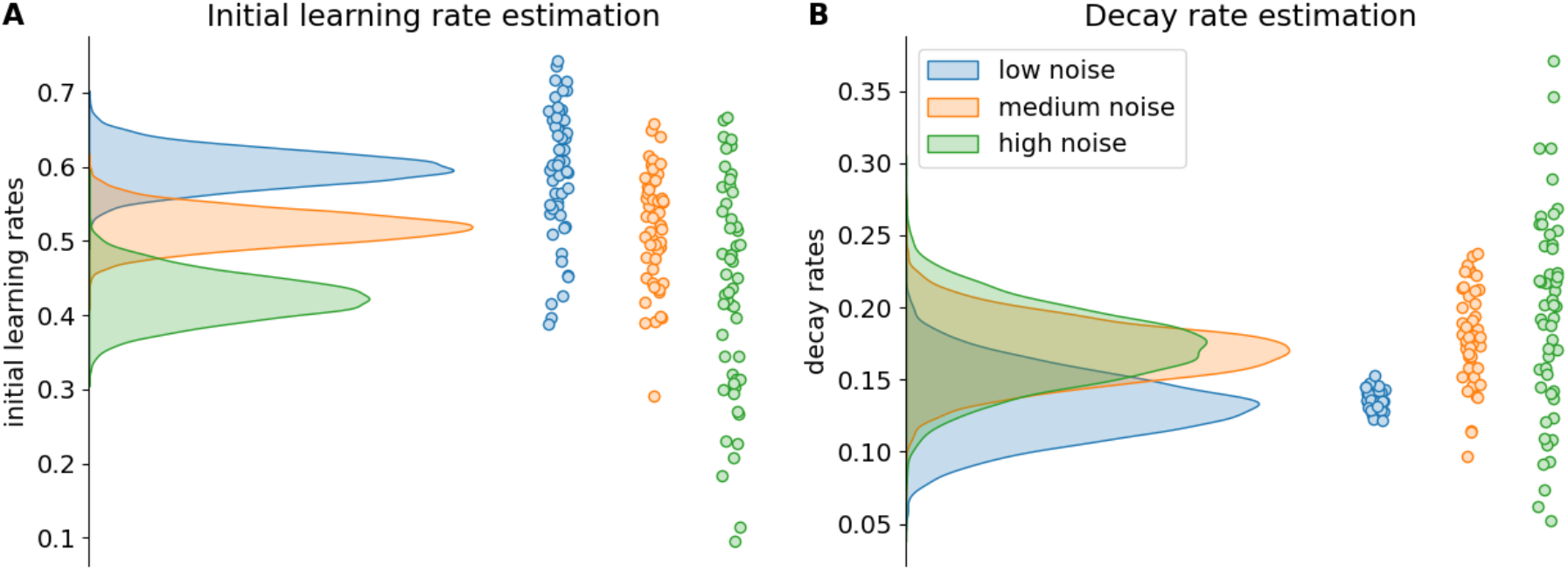
Bai model estimation results Experiment 2. The density plots on the left side of each subfigure show the full posterior densities over the means of the group-level distributions of the relevant parameters. The scatter plots on the right side of each subfigure show the means of all individual-level posterior distributions of the relevant parameters.

#### fMRI results

##### The neural representation of environment-specific learning rates during island presentation

To investigate the neural representation of environment-specific learning rates, we first performed a whole-brain searchlight representational similarity analysis (RSA) that tested for higher similarities between locations that required the same learning rate versus locations that required different learning rates. This analysis was done on data at the time of island presentation at the start of each block, which allowed us to test whether the mere presentation of the boat informing participants of where they would be fishing for crabs next triggered a state representation that differed depending on the relevant (initial) learning rate. Our hypothesis of representations that were specific to the high, mid, and low noise environments, but not for exact location, was encoded in a corresponding learning rate model representational dissimilarity matrix (RDM; Figure 6B). We then correlated this learning rate RDM with the corresponding neural RDM throughout the whole brain for each participant and we tested which voxels showed a significant (FDR-corrected *p* < .05) correlation on the group-level. This resulted in multiple clusters of significant voxels in left as well as right OFC (Figure 6C), in accordance with our hypotheses. We also found a large cluster of significant voxels in the occipital cortex. In the next paragraph, we further interpret the activation in the two clusters.

**Figure 6.**
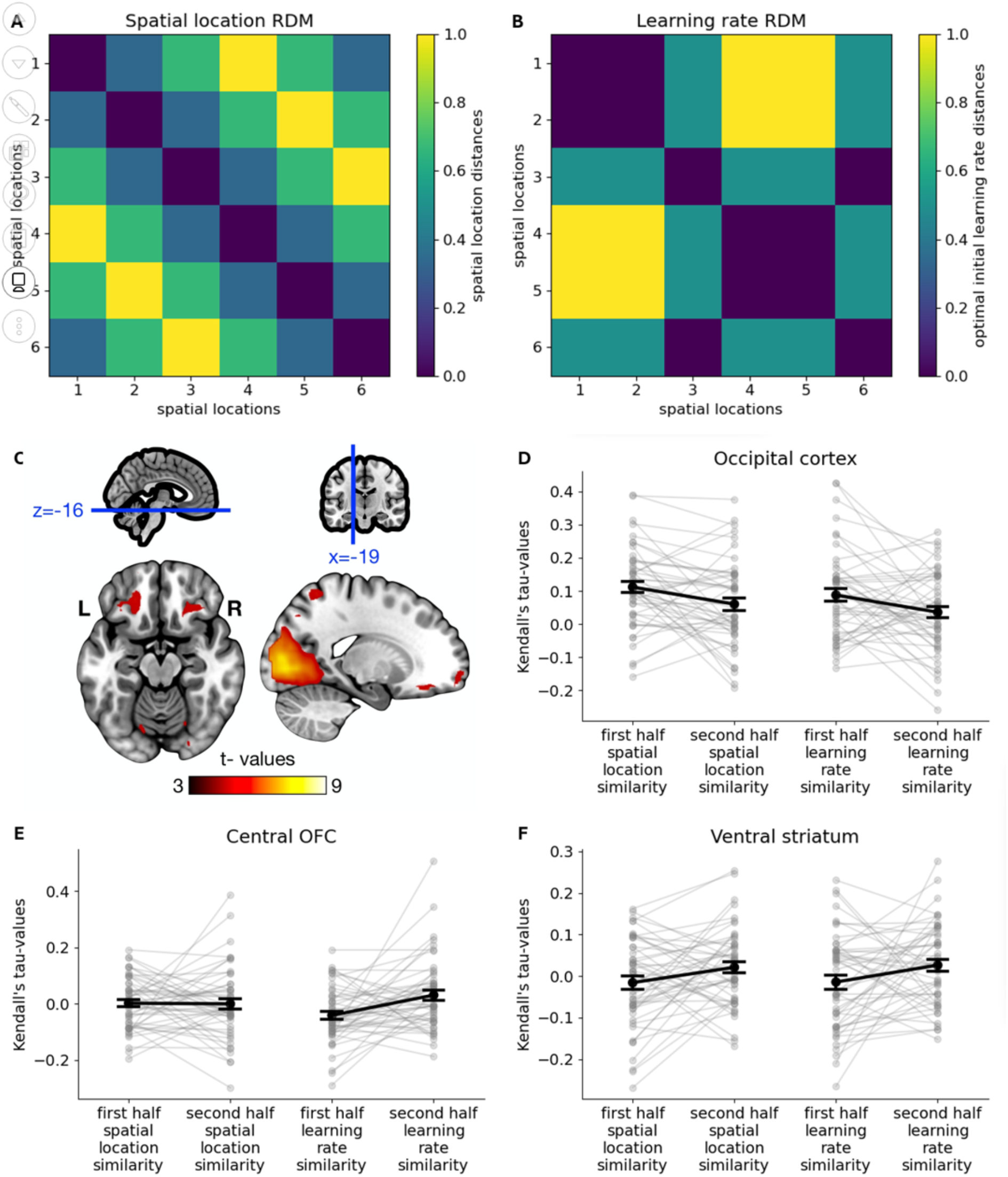
RSA analysis of the fMRI data. **A:** spatial location RDM. **B:** Learning rate RDM. **C:** Brain map of significant t-values resulting from the whole-brain searchlight RSA of fMRI data acquired while participants had just been transported to the next location around the island (correlation with learning rate RDM). **D-F:** Interaction effect between time (first vs. second half of task) and RDM (spatial location vs. learning rate RDM) in the occipital cortex, defined as the cluster of significant voxels found in the aforementioned whole-brain searchlight RSA (D); the central OFC as defined by (Kahnt et al., 2012), based on connections to other brain regions (E); and the ventral striatum, defined as the left and right nucleus accumbens according to the AAL atlas (F). Grey dots represent individual-level Kendall’s tau-values, while black dots and error bars represent group-level means and SEs of the means, respectively.

##### Dissociating the representation of spatial location and learning rate during island presentation

Since differences in optimal initial learning rate and differences in spatial location between the six locations are correlated, the activity in the occipital cortex is likely driven by spatial location rather than by learned initial learning rate. Crucially, representations of initial learning rate will tend to increase over blocks because they must be learned. Indeed, the behavioural data showed that environment-specificity of initial learning rates increased over time and was most pronounced in the second half of the task. Instead, representations of spatial locations should not. This allowed us to disentangle representations of spatial location and representations of initial learning rate. We thus performed a region of interest (ROI)-based follow-up analysis within the occipital cortex (defined as the cluster of significant voxels from the whole-brain RSA). For this analysis we constructed a second model RDM that encoded the different spatial locations, rather than the environments (spatial location model RDM; Figure 6A). We then calculated how strongly each participant’s neural RDM correlated with both model RDMs in the first half and the second half of the task separately. Finally, we performed a two (model RDM: spatial location vs. learning rate) by two (time: first vs. second half of the task) repeated measures ANOVA on the resulting correlations. We found a significant main effect of model RDM (*F*(1, 48) = 4.732, *p* = .035; Figure 6D), indicating that activity in the occipital cortex was indeed mostly driven by spatial location rather than initial learning rate, as well as a significant main effect of time (*F*(1, 48) = 6.665, *p* = .013), indicating that the representation of spatial location as well as initial learning rate decreased over time. We found no interaction effect between RDM and time.

To confirm that activity in the OFC was driven by differences in initial learning rate between the six locations and to pin down where exactly in the OFC environment-specific initial learning rates were represented, we divided the OFC into six subregions as defined by (Kahnt et al., 2012), based on its connections to other brain regions. We opted for this independent ROI approach, because the significant whole-brain cluster we observed only partially overlapped with OFC, as well as with other neighbouring regions. For each of these ROIs, we then calculated how strongly each participant’s neural RDM correlated with their learning rate RDM and with their spatial location RDM in the first half and the second half of the task separately. Finally, we performed a two (RDM: spatial location vs. learning rate) by two (time: first vs. second half of the task) repeated measures ANOVA on the resulting correlations. We found no main effect of RDM nor time in any of the OFC subregions, but we did find a significant interaction between RDM and time in one of the OFC subregions, namely the central OFC (*F*(1, 48) = 13.076, *p* < .001; Bonferroni corrected; Figure 6E), which is the OFC subregion that overlapped most with the largest cluster of significant voxels observed in the whole-brain RSA. Follow-up one-tailed paired t-tests confirmed that in the central OFC correlations between learning rate RDMs and neural RDMs were higher in the second half than in the first half of the task (*t*(48) = 3.051, *p* = .002), suggesting that representations of environment-specific initial learning rates increased over time. Instead, the correlation between spatial location RDMs and neural RDMs did not change across time (*t*(48) = -0.109, *p* = .543).

We also performed the same interaction analysis in the ventral striatum to study whether the ventral striatum also represented environment-specific initial learning rates during island presentation. We found no interaction effect between RDM and time, nor a main effect of RDM. However, we did find a main effect of time (*F*(1, 48) = 5.499, *p* = .023; Figure 6F), indicating that in the ventral striatum the representations of spatial location as well as initial learning rate increased over time.

##### Environment-sensitive neural processing of prediction errors during crab fishing

Finally, we also investigated how participants processed reward prediction errors. That is, we used the distance between the cage location (their prediction) and target location (the reward) as an approximation of reward prediction errors. To investigate neural activity during this phase we focused on the neural response to these prediction errors after the first and second trial, which was included as a parametric modulator in a series of first-level general linear models (GLMs). As a first analysis, we performed a whole brain (univariate) analysis on this parametric modulator, averaged over all environments, to evaluate whether we observed a typical response to prediction errors in the brain (Figure 7A-B). Indeed, in line with previous studies on reward prediction error processing (Calderon et al., 2021; O’Doherty et al., 2004; Pessiglione et al., 2006; Schultz et al., 1997), we observed significant clusters of voxels in the left and the right striatum, including in the ventral striatum.

**Figure 7.**
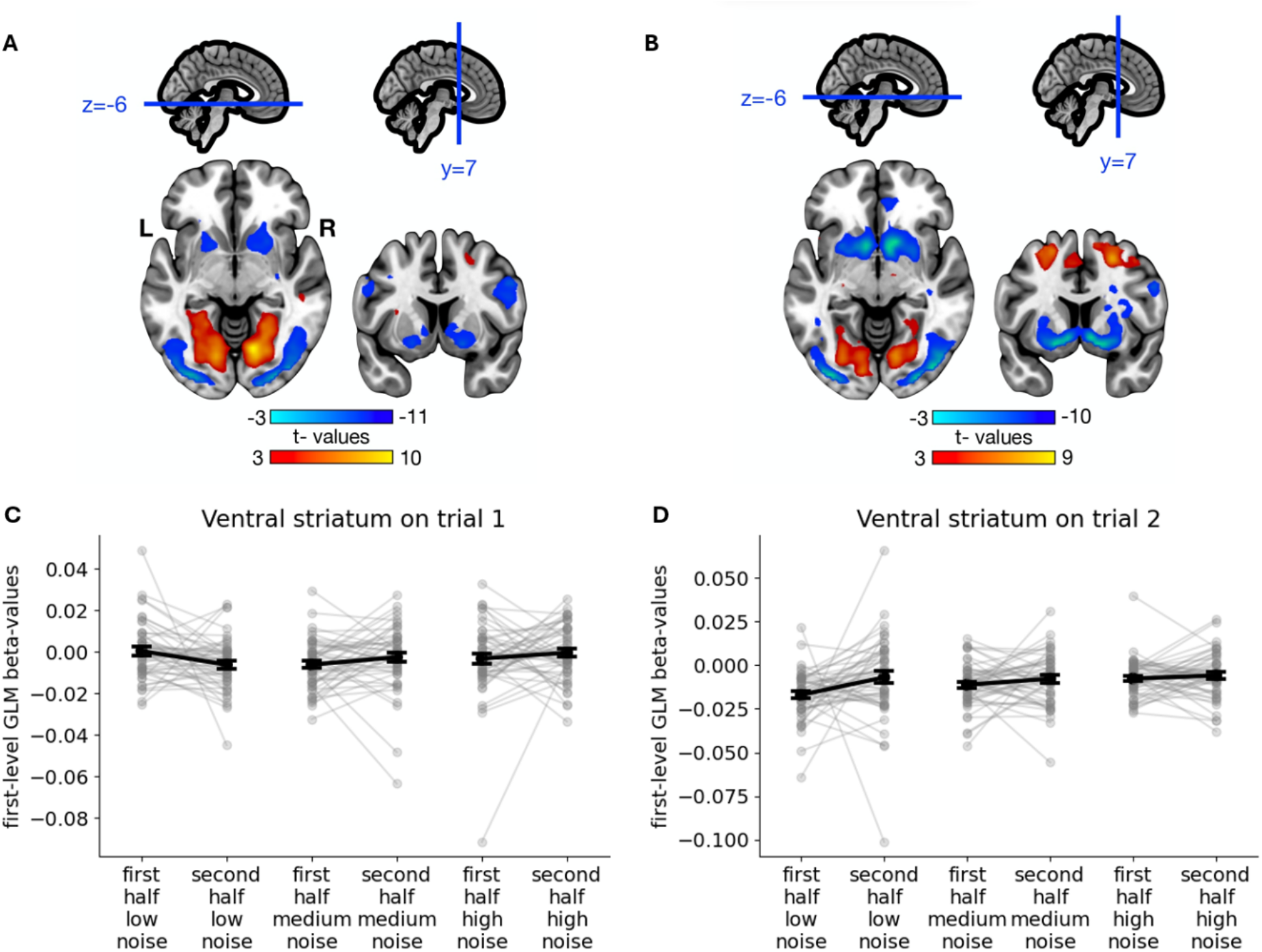
Results of the analyses of the effect of prediction error on the fMRI data. **A-B:** Brain map of significant t-values resulting from the whole-brain (univariate) tests of voxel activity being (parametrically) modulated by prediction error on trial 1 (A) and trial 2 (B). **C-D:** Interaction effect between time (first vs. second half of task) and environment (low vs. medium vs. high measurement noise) on the modulating effect of prediction error on trial 1 (C) and trial 2 (D) on ventral striatum activity. This ROI was defined as the left and right nucleus accumbens according to the AAL atlas. Grey dots represent individual-level GLM beta-values, while black dots and error bars represent group-level means and SEs of the means, respectively.

Next, we investigated whether there were interactions between environment (low vs. medium vs. high noise environment) and time (first vs. second half of the task) on these neural prediction error signals. Importantly, as mentioned above, our measure is only an (inverse) approximation of reward prediction error, because it measures the mismatch in location but not how much uncertainty people had around their point estimation. For example, seeing crabs appear on the exact cage location could still come with varying degrees of positive surprise depending on the uncertainty around their prediction, making it also difficult to estimate where exactly the prediction error turned form negative to positive. However, a region that is sensitive to the reward information in this signal, should show a neural response that is further dependent on environment and time, reflecting people’s ability to learn over time the more reward-informative nature of this signal in low noise environments. While we anticipated that the ventral striatum’s response would be modulated by the learning context, it is important to note that we held no specific a priori hypotheses regarding the direction of these effects.

We investigated this in the six OFC subregions and the ventral striatum using repeated measures ANOVAs. There were no significant effects in any of the OFC subregions. The ventral striatum, however, showed a significant interaction effect between environment and time (*F*(1, 48) = 4.222, *p* = .017; Figure 7C) in an analysis focusing on feedback processing during the first trial. Follow-up t-tests confirmed that the ventral striatum responded differently to prediction errors in the low noise environment in the second half compared to the first half of the task (*t*(48) = 2.284, *p* = .027), while it did not show such an evolution in the medium noise environment (*t*(48) = -1.468, *p* = .149), nor the high noise environment (*t*(48) = -0.842, *p* = .404). Specifically, ventral striatum activity showed a more negative response to larger location prediction errors. This pattern is consistent with its documented role in encoding reward prediction errors (Calderon et al., 2021). Namely, being closer to the centre of where the crabs appeared (i.e. smaller location prediction errors) corresponds to outcomes that are less negative or more positive than expected (i.e. smaller negative or larger positive reward prediction errors). On the second trial, there were no significant effects in any of the OFC subregions, while the ventral striatum only showed a significant main effect of time (*F*(1, 48) = 4.884, *p* = .032; Figure 7D), suggesting it may have become less sensitive to reward prediction errors on the second trial over time.

## Discussion

Humans can adapt how they learn in response to environmental demands, but how these adaptations unfold over time, and how to dissociate different types of adjustments, remains poorly understood. Here, we developed a new paradigm that systematically disentangles two types of learning rate adaptations: a fast and transient response to local prediction errors, and a slower form of meta-learning an optimal learning rate as a function of higher-level environmental statistics. Specifically, we designed a gamified task in which participants fish for crabs on six different fishing locations around an island. The environmental statistics about the crabs’ hiding spots implied different optimal learning rates for the different locations. We extracted participants’ learning rates on each trial, which allowed us to test both (1) whether participants dynamically adjusted their learning rates in response to locally experienced prediction errors and (2) whether we could observe, above and beyond these local adaptations, different initial learning rates tailored to the environmental statistics. Across two experiments, we observed that participants did both: They immediately adapted their learning rates on a trial-by-trial basis in response to just-experienced prediction errors, but also learned, over time, to use different initial learning rates on the different locations around the island.

Computational modelling confirmed our findings. We fitted six models to the data, and in both experiments the best fitting model was the environment-specific Bai model, which implemented both the learning of environment-specific initial learning rates (across blocks) and learning rate updating proportional to recently experienced prediction errors (within blocks). While the Kalman filter provides optimal learning rates for the present task on every trial based on underlying environment statistics, the Bai model assumes a (learned) initial learning rate which is up- or downregulated by experienced prediction errors. According to the Kalman filter, learning rates should quickly decrease towards 0 irrespective of the initial learning rate, and independently of (individual differences in sensitivity to) experienced prediction errors. While participants’ learning rates do decrease over trials, they stabilise around 0.3, indicating that people are more responsive to noise than is optimal.

Overall, our behavioural data and modelling suggest that, over the course of both experiments, participants learned to instantaneously retrieve relevant learning rates upon arrival on any given location around the island. Our findings go beyond previous studies that documented changes in learning rates over time (Behrens et al., 2007; Browning et al., 2015; Goris et al., 2021), by showing that people can also switch back and forth between learning rates across environments (see also (Simoens et al., 2024)). As such, the present study provides support for recent theories of meta-learning (Botvinick et al., 2009; Holroyd & Verguts, 2021; Silvetti et al., 2018; Wang et al., 2018), as well as cognitive control (Abrahamse et al., 2016; Braem et al., 2019; Chiu & Egner, 2017), which posit that cognitive control is implemented as the environment-specific regulation of task execution parameters, such as learning rate. Interestingly, our work is also consistent with that of others who have shown that also the local adaptation strategy in itself, may reflect an environment-specific strategy based on prior experience with these environments’ generative structure (Bakst & McGuire, 2021, 2021; Lee et al., 2020).

We also investigated which brain regions represent sustained, meta-learned associations between environments and learning rates, enabling one to instantaneously retrieve the relevant learning rate when revisiting an environment. Importantly, previous studies examined neural correlates of learning rates during outcome evaluation, where learning rates may be adjusted online as a function of locally experienced prediction errors (e.g., (Behrens et al., 2007; Browning et al., 2015; Nassar et al., 2012). In contrast, our RSA analysis targeted neural activity at island presentation, before any outcome information was available. At this moment, learning rates cannot be updated based on current feedback and instead reflected the retrieval of a previously learned, environment-specific learning-rate settings. This difference reflects our hypothesis that the OFC represents the latent states in a cognitive map of the task (Knudsen & Wallis, 2022; Moneta et al., 2024; Schuck et al., 2018; Wilson et al., 2014), which are expected to activate as soon as the agents can infer which task state it is in. Several studies have identified such “partially observable” task states in the medial OFC (Bradfield et al., 2015; Schuck et al., 2016; Tan et al., 2025; Wimmer & Büchel, 2019), in line with the region identified here (but see e.g., (Ongur & Price, 2000), for important anatomical distinctions between medial and lateral OFC and (Tan et al., 2025) for an example of related functions in lateral OFC). Our finding extends this notion by suggesting a link between OFC and meta learning, wherein meta-learned information becomes encapsulated in task states (Hattori et al., 2023; Moneta et al., 2024). Consistently, OFC has been shown to represent task states (Moneta et al., 2024; Stalnaker et al., 2015; Wilson et al., 2014). While earlier evidence shows that the OFC represents concrete aspects of task states, such as task-relevant stimulus features (Schuck et al., 2016), we hypothesized that the OFC also represents more abstract aspects, such as learned, environment-specific learning rates. Indeed, we showed that the central OFC gradually came to represent these environment-specific learning rates (or the environment-specific statistics that drive them). While previous work speculated that these different levels could have different neural underpinnings (Sharpe et al., 2019), our findings indicate OFC might signal states on multiple levels. This does not imply identical learning dynamics; fast-changing trial-specific states might be learned through activity dynamics, while higher-level contextual states could involve synaptic plasticity.

We further observed that the ventral striatum learned to differentially respond more to positive reward prediction errors (or less to negative reward prediction errors) depending on the currently relevant environment-specific learning rate. Similar to previous studies, the ventral striatum responded more to prediction errors where the noise was lowest (Diederen et al., 2016) but see (Mah et al., 2024). This heightened sensitivity of the ventral striatum to reward prediction errors in low-noise environments aligns with the fact that these signals are most behaviorally informative in stable contexts, where a higher learning rate is optimal for guiding future behavior. The ventral striatum also became less sensitive to the second trial’s prediction error over time. Presumably, as participants gained more experience with the task’s global reward structure (specifically that all targets center around a fixed mean), the first reward prediction error per round became the primary source of information, rendering subsequent reward prediction errors within that round exponentially less behaviorally relevant over time. We also saw a prediction error signal in the ACC, but only on the second trial. Interestingly, prediction errors on the second trial are the first events after participants have formed an initial, local estimate of the crab’s location and prediction errors could meaningfully signal the need to update internal models or control settings (Hayden et al., 2011; Silvetti et al., 2018), in line with earlier studies suggesting the ACC’s role in uncertainty-driven belief updating in this task (McGuire et al., 2014). Taken together, our fMRI data analyses suggest that the OFC, more specifically the central OFC, represents environment-specific learning rates, which may in turn affect how the ventral striatum and ACC responded to prediction errors.

Our findings are in line with recent conceptualizations of learning across two time scales, both in artificial and biological (Binz et al., 2024) agents. Here, fast learning would presumably occur in (neural) activation space, whereas slower learning would occur in weight space. Although our study did not allow specifying the locus of (fast and slow) learning, a direct comparison between artificial and biological agents was reported in (Hattori et al., 2023). They used a two-armed bandit task to study how mice as well as deep reinforcement learning models adapt their behaviour over time. They found that, in mice, the OFC is crucial for slow, across-session learning, which gradually refines neural circuits that support fast, within-session learning. The researchers also observed parallels between the mechanisms underlying learning at these two different time scales in mice and deep reinforcement learning models. That is, both systems employed synaptic plasticity mechanisms to shape neural connectivity for learning on the slow time scale, but instead on recurrent activity dynamics for learning on the fast time scale (Duan et al., 2017; Wang et al., 2018). Indeed, blocking synaptic plasticity in the OFC disrupted across-session learning, but left within-session learning intact in expert mice (Hattori et al., 2023). Thus, slow, across-session meta-learning may involve plasticity-based mechanisms in the OFC, which serve to improve fast, within-session learning of new tasks through recurrent activity dynamics. Here, we provided first evidence that a similar dual process may operate in humans.

A recent theme in artificial intelligence and computational neuroscience is that (artificial and biological) agents (should) learn to cluster the environments they are confronted with; and associate different (low- or high-level) parameters to each such environment (Collins & Frank, 2013; Verbelen et al., 2022). This approach leads to efficient learning, not least because it shields against catastrophic interference. Here, we used only three environments (albeit on six locations), so the clustering was relatively easy in our case. However, future studies could use a design similar to the one presented here to investigate the neural underpinnings of clustering environments (and associated learning rates) in a continuous range of environments around the island. Similarly, the present study focused on differentiating between environments in terms of learning rate. However, human as well as nonhuman agents are often confronted with novel situations, in which they may instantaneously deduce appropriate settings for task execution parameters such as learning rate, across contexts; in brief, adaptive agents can generalize task execution parameters across similar contexts. Future studies could leverage the task developed for the present study to investigate the neural underpinnings of this generalisation of (abstract) knowledge by, for example, introducing more locations later on in the task.

In conclusion, the present study demonstrates the importance of differentiating between two time scales of adaptation in human learning rates. Fast time scale learning within environments and slow time scale learning about environments likely involve different cognitive processes with different neural underpinnings. Nevertheless, this distinction has thus far been largely overlooked, in favour of the fast time scale. Future research could leverage the experimental design presented here to further investigate the slow time scale as well. Our gamified design (Allen et al., 2024) can also provide new ways to test theories about the relation between meta-learning and development (Nussenbaum & Hartley, 2024) or psychological pathologies, such as theories of autism which posit that autism is related to deficits in the detection of environmental differences in learning opportunities (Goris et al., 2021; van de Cruys et al., 2014).

## Methods

### Participants

50 participants (42 female, 8 male) were recruited for Experiment 1, and 53 participants (41 female, 12 male) for Experiment 2, through Sona (https://www.ugent.sona-systems.com/). All participants were between 18 and 35 years old. Because all participants caught about the same number of crabs, no participants were excluded from the behavioural data analyses. Since no response time deadline was implemented, no trials were excluded from the analyses. However, two participants were excluded from the fMRI data analyses because they were left handed, and another two because of technical problems with the fMRI data acquisition.

Experiment 1 was approved by the Ghent University Psychology and Educational Sciences Ethical Committee, and Experiment 2 by the Ghent University Medical Ethical Committee. Participants signed informed consents prior to participation. Experiment 1 took participants about 45 minutes to complete, in return for which they received a participation fee of €10. Participants in Experiment 2 received a participation fee of €35, because we also administered fMRI recording. In both experiments, the participant who caught the most crabs, received a €50 gift certificate for bol.com (an online store that offers general merchandising products). All data and code can be found on https://osf.io/qft2p for Experiment 1, and on https://osf.io/be4td for Experiment 2.

### Experimental design

In Experiment 1, participants performed a novel crab-fishing task, which was programmed in jsPsych (de Leeuw et al., 2023). At the start of each of 60 blocks, a boat took them to one of six locations around an island (Figure 1C). There they dropped a cage ten times (trials) trying to catch crabs. Each time a cage was dropped, one location was sampled from the sampling distribution 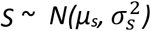, truncated between *µ*_*s*_ ± 1.65 * *σ*_*s*_ in order to avoid confusing participants with the occasional extreme outlier as well as off-screen locations. Before the cage reached the ocean floor, five crabs appeared and spread out evenly from this location, each of which was either caught by the cage or ran away. At the beginning of each block, *µ*_*s*_ was sampled from the prior distribution 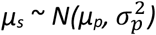, truncated between *µ*_*p*_ ± 1.65 * *σ*_*p*_, with *µ*_*p*_ = the centre of the screen.

Crucially, the standard deviations of both prior and sampling distributions were dependent on the location around the island. On two randomly selected adjacent locations, *σ*_*p*_ was large (18.75% of the screen width), while *σ*_*s*_ was small (6.25% of screen width), making this a low noise environment. Here, a high initial learning rate is optimal since the mean of the sampling distribution could be far away from the centre of the screen, but all crabs will cluster close together. Thus, in estimating the mean of the sampling distribution, it makes sense to give a lot of weight to the first crabs (i.e., use a high learning rate), and exponentially decrease the learning rate afterwards. On the two adjacent locations on the exact opposite side of the island, the situation was reversed, making this a high noise environment. Here, a low (initial) learning rate is optimal since the mean of the sampling distribution can only be near the centre of the screen, but crabs can appear far away from each other (and the centre of the screen). Thus, in estimating the mean of the sampling distribution, it makes sense not to give too much weight to any individual crabs (i.e., use a low learning rate). Finally, on the two locations in between, the standard deviations *σ*_*p*_ and *σ*_*s*_ were intermediate (12.5% of screen width) and equal to each other, making this a medium noise environment and requiring an intermediate (initial) learning rate.

Participants visited all six locations around the island once in randomised order before visiting all locations a second time in (re)randomised order, and so on. The sailing of the boat from the previous to the next location unfolded over three to five seconds, depending on how far apart the locations were (one second stationary at previous location, followed by 60 degrees per second of sailing around the island, followed by one second stationary at the next location). Next, participants could move the cage to the left using the f-key and to the right using the j-key. Tapping the key would move the cage 1% of the screen width in the corresponding direction, while holding the key would slide the cage in that direction more quickly. There was no response time deadline. Participants could drop the cage by pressing the space bar, after which feedback unfolded over the course of 1.5 seconds. For the first 500 ms the cage sank until halfway to the bottom of the ocean; for the next 500 ms the cage sank all the way to the bottom of the ocean while five crabs appeared out of one point on the ocean floor and spread out to cover the same proportion of the screen width as the cage (18.75%); for the last 500 ms crabs that were not caught by the cage ran away (see Figure 1C), while crabs that were caught by the cage remained in place. The usage of a wide cage as well as five crabs was to ensure that participants could still catch some crabs in the high noise environment. At the start of each trial after the first trial (within blocks), the cage was moved back to the top of the screen at the x-coordinate where it was dropped on the last trial. The heap of sand where five crabs had crawled out of the sand on the last trial, would still be visible in order to help participants determine where to drop the cage next. While participants were fishing for crabs, a radar at the top of the screen reminded them of where around the island they were at all times.

Participants were instructed that around this island, crabs live in groups that are denser near the centre than towards the edges and that the local group of crabs would not change location while they were fishing on its location. The local group of crabs would only change location while they were fishing somewhere else. Hence, they should try to drop their cage over the centre of the group to maximise reward. They were also instructed that these groups of crabs might have different sizes around the island, so that they should keep track of where around the island they are.

After receiving the instructions, participants first performed a short practice phase during which they performed one block in each of the three environments. During the practice phase, participants could not see where around the island they were, while the (normally distributed) group of crabs under the sand was made visible while they were fishing.

During the actual task, the group of crabs was, of course, not visible while participants were fishing for crabs. However, they did receive block feedback at the end of each block. That is, at the end of each block, the group of crabs was made visible, as well as all ten (heaps of sand at) locations where crabs had crawled out of the sand during the block. During the fishing, only the location where crabs had appeared after the previous cage was dropped, was made visible as a little heap of sand. Finally, the task was divided into four rounds, in between which participants could take a short break.

Experiment 2 was programmed in PsychoPy (Peirce, 2007) and consisted of 60 blocks that were identical to the blocks in Experiment 1, except that they consisted of only eight trials. Additionally, 60 blocks that consisted of only two trials were randomly intermixed with the 60 longer blocks. These blocks were included to increase power for analyses on the very first (i.e., the most informative) trials within blocks. At the end of these shorter blocks, participants received no block feedback.

To make the design suitable for the MR-scanner, some additional minor changes were made. At the start of a block, the boat did not sail to a new location around the island but was immediately presented at the new location for a duration of three to seven seconds. Next, participants could move the cage to the left and to the right using their right index finger and right middle finger, respectively, on a response box that was placed in their right hand. Participants could drop the cage using their left index finger on a response box placed in their left hand. A laser pointer was added to the cage that pointed straight down from the centre of the cage to the ocean floor. Feedback unfolded over the course of 750 ms. During the first 250 ms, the cage sank until halfway to the bottom of the ocean. During the next 250 ms, the cage sank further to the bottom of the ocean and five crabs appeared out of one point on the ocean floor and spread out to cover the same proportion of the screen width as the cage. Before the start of the last 250 ms, crabs that were not caught by the cage disappeared from the screen. After the first and second trial of each block, there was an intertrial interval of three to seven seconds during which only the heap of sand where crabs had just crawled out of the ocean floor as well as a red cross (also on the ocean floor) at the centre of where the cage had just been dropped remained visible on the screen. Finally, block feedback lasted for three to seven seconds.

### Behavioural data analysis

All data and analysis scripts can be found on the Open Science Framework. Participants’ continuous responses allowed us to solve the delta rule:

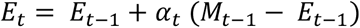

where *E*_*t*_ and *E*_*t*-1_ are the participant’s estimates of *µ*_*s*_ on trials *t* and *t* - 1, respectively (i.e., the locations where the participant dropped the cages), *α*_*t*_ is the participant’s learning rate on trial *t*, and *M*_*t*_ is the location where the five crabs appeared on trial *t*, for the learning rate *α* for each trial *t >* 1. That is, each trial *t* > 1 we calculated:

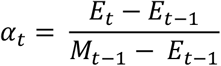

This allowed us to test whether participants showed a decrease in learning rate over the course of a block, as they learned more and more about the location of crabs within that block. To this end, we performed a linear mixed effects model analysis, with a random intercept for participant and a random slope for environment as well as for trial number (as well as for the environment-trial interaction), on participants’ median learning rates (for each environment-trial combination). We opted for median rather than mean learning rates (on a participant-level) because we observed some extreme outlier learning rates on trials directly following a trial in which crabs appeared directly next to where participants dropped their last cage. Here, any movement of the cage has an effect on the resulting learning rate which is unlikely to be proportional to participants’ actual learning rates (by blowing up the numerator, either positively or negatively, in the above formula for *α*_*t*_.

To investigate whether participants’ median learning rates on the second trial of each block were significantly different in the three environments and whether participants gradually learned to use different initial learning rates in the different environments, we divided Experiment 1 into two halves and Experiment 2 into four quarters (i.e., the four functional runs in the MR-scanner) and performed a two (time: the two halves) or four (time: the four runs) by three (environment: the three environments) repeated measures ANOVA on participants’ median learning rates on the second trial of each block. We interpreted the significant main effect of environment and the significant interaction effect between time and environment using one-tailed paired t-tests. We opted for median rather than mean learning rates (on a participant-level) because we observed some extreme outlier learning rates on trials directly following a trial in which crabs appeared directly next to where participants dropped their last cage. Here, any movement of the cage has an effect on the resulting learning rate which is unlikely to be proportional to participants’ actual learning rates (because the denominator in the formula for *α*_*t*_ is close to zero and the equation hence unstable).

### Model estimation and selection

We fitted six models to the data using hierarchical Bayesian analyses (HBA) (Ahn et al., 2017). The HBA was performed in Stan (Carpenter et al., 2017), which uses Hamiltonian Monte Carlo (HMC) sampling, a variant of Markov chain Monte Carlo (MCMC) sampling. The first two models were the environment-specific and non-environment specific versions of the Kalman filter. The Kalman filter assumes that people update their estimates using the delta rule, with a learning rate that depends on both estimate and measurement uncertainty:

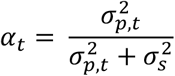

where *α*_*t*_ is the participant’s current learning rate; 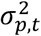 is the participant’s current estimate uncertainty, which initially is the variance of the prior distribution; and 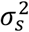 is the measurement uncertainty, which here is the variance of the sampling distribution. Crucially, learning rate decreases over time because:

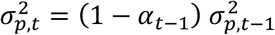

Initial estimate uncertainty 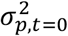 and measurement uncertainty 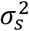 cannot both be free parameters, because they trade off against each other, which would result in bimodal and unreliable posteriors (Daw et al., 2006). Therefore, we fixed measurement uncertainty to 0.125^2^ (the true value in the medium noise environment) and only estimated initial estimate uncertainty. As an alternative procedure, we also tried fixing measurement uncertainty to the true value in each environment separately. The two approaches resulted in different posterior densities for estimate uncertainties, but in similar posterior densities for learning rates. Moreover, both approaches resulted in almost identical values for the LOOIC. We conclude that we can safely fix measurement uncertainty to 0.125^2^ across environments. In the environment-specific version of the model, each participant was assumed to use separate initial estimate uncertainties for each environment, while in the non-environment-specific version of the model, participants were assumed to use the same initial estimate uncertainty in both environments.

The next two models were the environment-specific and non-environment specific versions of the Rescorla-Wagner model, which assumes that people update their estimates according to the delta rule (with a constant learning rate). In the environment-specific version of the model, each participant was assumed to use separate learning rates for each environment, while in the non-environment-specific version of the model, participants were assumed to use the same learning rate in both environments.

The final two models were the environment-specific and non-environment specific versions of the Bai model (Bai et al., 2014), which is in the general family of models that adapt learning rate as a function of prediction errors (Krugel et al., 2009; Pearce & Hall, 1980). Hence, the Bai model also supposes that people update their estimates according to the delta rule, but use a learning rate dependent on recently experienced prediction errors:

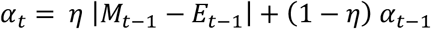

where *η* is a decay rate, which also takes a value between 0 and 1 and determines how much learning rates are affected by recently experienced prediction errors. That is, when prediction errors are large, learning rate increases, but when prediction errors are small, learning rate decreases exponentially. In the environment-specific version of the model, each participant was assumed to use separate initial learning rates and decay rates for each environment, while in the non-environment-specific version of the model, participants were assumed to use the same learning rate and decay rate in both environments.

For each environment (low vs. medium vs. high noise), individual-level free parameters were assumed to be drawn from a group-level normal distribution specific to that environment. For the means of these group-level distributions, we used uniform priors between 0 and 1, while for the standard deviations we used half-Cauchy (0, 5) priors. For the individual-level parameters we also used bounded uniform priors. To minimize the dependence between the means and standard deviations of group-level distributions, we used non-centred parameterisations. To maximise the efficiency of HMC sampling, parameters were first estimated in an unbounded space and then probit-transformed to the relevant bounded space (Ahn et al., 2017).

For each model, 4000 samples were drawn from the posterior distributions, the first 1000 of which were discarded as burn-in, across four sampling chains, resulting in a total of 12,000 posterior samples. Convergence of posterior distributions was checked by visually inspecting the traces and by numerically checking the Gelman-Rubin statistics (Gelman & Rubin, 1992), which were all well below 1.1, for each estimated parameter.

We used the LOOIC (Vehtari et al., 2017) for model comparisons. The LOOIC approximates the log pointwise posterior predictive density of observed data, which is the out-of-sample predictive accuracy of a model. Therefore, higher values indicate higher out-of-sample predictive fit of the model to the data.

To calculate the posterior probability that learning rates were higher in one environment than in another, we calculated the proportion of posterior samples in which the group-level mean learning rate was higher in the one environment than in the other. Thus, a posterior probability higher than 95% corresponds to a one-tailed p-value lower than 0.05.

### Parameter recovery

To make sure that the experimental design and model estimation procedure would allow for the reliable estimation of the model parameters, we performed parameter recovery simulations. Specifically, using the aforementioned design details, we simulated 16 datasets with each of the three models in each of the three environments separately. For the RW model we used each of the learning rates {0.2, 0.4, 0.6, 0.8} four times as the true mean of the group-level distribution of this parameter. For the Kalman filter we used each of the estimate uncertainties {0.25, 0.5, 0.75, 1} four times as the true mean of the group-level distribution of this parameter. For the Bai model we used each combination of learning rates {0.2, 0.4, 0.6, 0.8} and decay rates {0.2, 0.4, 0.6, 0.8} as the true mean of the group-level distribution of these parameters. For each parameter, we then randomly sampled 50 true parameter values from the group-level normal distribution with the relevant mean and an SD of 0.2 for learning rates and decay rates, and 0.25 for estimate uncertainties. We then simulated a dataset using these parameter values and the experimental design described above and fitted the model to this simulated dataset using our model estimation procedure. Finally, for each model and each environment separately, we calculated parameter recovery rates by correlating all (16 datasets x 50 participants = 800) true parameters to the corresponding estimated parameters, that is, the individual-level posterior means. These simulations indicated that the model could be reliably fitted to our behavioural data. For learning rates in the RW model, estimate uncertainties in the Kalman filter, and initial learning rates in the Bai model recovery rates were higher than 0.975 in each environment. For decay rates in the Bai model parameter recovery rates were 0.807, 0.868 and 0.918 for the low, medium and high noise environment, respectively.

### Model recovery

To ensure that the experimental design and model selection procedure would allow for the reliable selection of the model fitting the data best, we also performed model recovery simulations. Specifically, using the experimental design described above, we simulated 10 datasets with each of the six models described above. For each model parameter, we used the estimated posterior mean and SD of the group-level distribution (of the empirical data) as the mean and SD, respectively, of the group-level distribution (of the simulated data), and randomly sampled 50 values from this distribution. For each model, we then simulated a dataset using these parameter values and the experiment design described above and fitted all six models to this simulated dataset using the model estimation procedure described above. Finally, using the model selection procedure described above, we tested for each simulated dataset which model fitted it best and calculated model recovery rates as the proportion of datasets simulated with each of the three models that was best fit by the correct model. Model recovery rates were 100% for each of the six models.

### Model validation

For validating the winning model, we used the posterior predictive check method (Gelman et al., 1996). This method takes participants’ fitted model parameters and uses them to simulate responses given their individual trial sequence. Simulated and true response patterns can then be compared to determine how well the model captures participants’ behaviour (Palminteri et al., 2017). Specifically, for each participant, we used the means of the participant-level posterior densities over the relevant parameters to simulate a response sequence conditional on the trial sequence this participant had received. We then (Pearson) correlated each participant’s true response sequence with the simulated response sequence.

Posterior predictive checks indicated that the environment-specific Bai model indeed adequately captured participants’ behaviour. The average correlation between participants’ true choice sequences and the response sequences obtained by simulating data using their parameter estimates was 0.793 in Experiment 1 and 0.863 in Experiment 2. More specifically, the average correlations were 0.927, 0.81 and 0.642 for the low, medium and high noise environment, respectively, in Experiment 1 and 0.955, 0.879 and 0.754 for the low, medium and high noise environment, respectively, in Experiment 2. Although still medium to large and reliable, we do note that these correlations were slightly lower in the high noise environment. In this environment, individual outcomes are considerably less indicative of the latent mean, which may reduce the usefulness of the trial-by-trial, prediction-error–driven learning-rate adjustments that we see in the low and medium noise environments. Under extreme conditions of variability, people may rely less on delta-rule updating and more on alternative strategies (d’Acremont & Bossaerts, 2016; Reynders et al., 2026), such as exploratory adjustments or heuristics that are not explicitly captured by the Bai model, but also outside the scope of the present paper.

### fMRI data acquisition and preprocessing

T1-weighted MPRAGE structural images (1 mm isotropic voxels, 256 x 256 matrix, 176 axial slices, 9° flip angle), GRE field map images (528 ms TR, 7.38 TE, 60° flip angle), and T2*-weighted EPI functional data (2.5 mm isotropic voxels, 64 x 64 matrix, 1780 ms TR, 27 ms TE, 66° flip angle) were acquired on a 3 T Prisma scanner system (Siemens) with a 64 channel head coil. Functional data were acquired in 4 runs, each of which lasted about 12 minutes.

MRI data were preprocessed using fMRIPrep 23.1.0 (Esteban et al., 2019) and involved motion correction, field map based geometric undistortion, slice timing correction, coregistration of anatomical and functional scans, normalization into MNI space, and spatial smoothing with a 5 mm FWHM Gaussian kernel.

### fMRI data analyses

First-level (subject-wise) general linear modelling was done in SPM (https://www.fil.ion.ucl.ac.uk/spm/) and involved regressors of interest that captured stimulus onset events (using an event-related design (i.e., with all event durations = 0); see below) and nuisance regressors that reflected participant movement (six regressors) as well as the global signal (one regressor). All regressors were convolved with a canonical hemodynamic response function.

Standard GLMs were used to estimate voxel activations associated with stimulus display. First-level models included separate regressors for each of the seven events of interest in each experimental block crossed with each of the six locations around the island plus the above described seven nuisance regressors for each run separately. The seven events of interest (regressor onset times) were: (1) presentation of the island at the beginning of each block, (2) the onset of the first trial, (3) the end of the feedback of the first trial, (4) the onset of the second trial, (5) the end of the feedback of the second trial, (6) the onsets of each remaining trial in the experimental block (when it was a long block), and (7) the onset of the block feedback (when it was a long block). We also included (mean-centred absolute) prediction error (in the current trial) as a parametric modulator for events 3 and 5 separately (as well as for each location around the island separately). This resulted in 244 whole brain maps of parameter estimates (“betas”; ((seven events of interest + two parametric modulators) * six locations around the island + seven nuisance regressors) * four runs).

The RSA focused on the beta maps capturing voxel activation during the presentation of the island at the beginning of each block. Before proceeding to the RSA, we normalized each voxel to its own within-run mean, by subtracting the voxel’s overall mean activation from each location’s activation, so that the overall mean is zero (Diedrichsen & Kriegeskorte, 2017). Next, we performed a whole-brain searchlight RSA in three steps. Firstly, for each participant, we constructed a 24×24 design-based RDM with as rows and columns the six locations in each run (out of 4) and in each cell the dissimilarity between the optimal initial learning rate in the row’s location and the optimal initial learning rate in the column’s location (which could be 0, 1, or 2). Secondly, for each participant, we went through the entire brain in spherical searchlights with a radius of three voxels to construct 24×24 neural RDMs and compare (the upper triangle of) these RDMs to (the upper triangle of) the design-based RDM (also excluding within run cells) by computing Kendall’s Tau. Within each searchlight, the neural dissimilarity between each pair of locations was computed as one minus the Pearson correlation between the voxel wise activations for those locations. This resulted in a brain map of dissimilarity values (tau-values) for each participant where each voxel’s tau-value is computed with that voxel in the centre of a searchlight. Finally, we used one-tailed t-tests to test which voxels had tau-values significantly higher than zero, which indicates that they responded differently to different locations around the island, using a significance threshold of 0.05 corrected for multiple comparisons using the false discovery rate (FDR).

Next, we performed a ROI-based follow-up analysis with the occipital cortex, defined as the cluster of significant voxels from the whole brain searchlight RSA, as ROI in order to tease apart representations of spatial location and initial learning rate in this ROI, which were strongly correlated with each other. For this analysis we constructed a second 24×24 design-based RDM with as rows and columns the six locations in each of the four runs, and in each cell the dissimilarity between the spatial location in the row’s location and the spatial location in the column’s location (which could be 0, 1, 2, or 3). We then correlated the parts of (the upper triangle of) both design-based RDMs that concerned the first two runs (excluding within run cells) to the corresponding part of the neural RDMs, and then did the same for the last two runs. Finally, we performed a two (design feature: spatial location vs. optimal initial learning rate) by two (time: first vs. second half of the task) repeated measures ANOVA on the resulting correlations.

Next, we performed more in depth RSAs within predefined anatomical ROIs within the OFC to test whether within these regions the representation of initial learning rate level gradually became stronger than the representation of spatial location. To this end, we performed four RSAs within each ROI. That is, we correlated the parts of (the upper triangle of) both design-based RDMs that concerned the first two runs (excluding within run cells) to the corresponding part of the neural RDMs, and then did the same for the last two runs. Finally, we performed a two (design feature: spatial location vs. optimal initial learning rate) by two (time: first vs. second half of the task) repeated measures ANOVA on the resulting correlations. Where a significant interaction effect was found, we used one-tailed paired t-tests to interpret this interaction effect. The ROIs were created using SPM’s wfupickatlas toolbox. The OFC subregions were defined as in, based on connections to other brain regions. Here, we defined multiple ROIs within the OFC based on what is known about its structure, rather than defining an ROI based on the cluster of significant voxels observed in the whole brain RSA described above, as we did for the occipital cortex, because we wanted to check whether there were no significant effects in ROIs in which clusters of voxels did not exceed the stringent significance threshold applied in the whole brain RSA described above. Since we tested six ROIs, we used Bonferroni correction for multiple comparisons, which lowered the significance threshold to .05/6 = .008.

Although we found no significant clusters of voxels in the ventral striatum in the whole-brain RSA described above, we also performed the same repeated measures ANOVA we performed in the occipital cortex and the OFC in the ventral striatum to check if there was no effect there that did not survive the stringent significance threshold applied in the whole-brain RSA (or was cancelled out by an interaction between time and RDM, considering the whole-brain RSA exclusively tests for a main effect of RDM). This ROI was created using SPM’s wfupickatlas toolbox and was defined as the left and right nucleus accumbens according to the AAL atlas.

We also performed all of the (whole-brain and ROI-based) RSAs described above on neural data acquired during the presentation of feedback after the first two trials in each block.

Finally, we also checked the effect of prediction errors on brain activity using univariate analyses. First, as a sanity check, we used a whole-brain approach in which we averaged together (over blocks and environments) all whole-brain maps of first level beta-values for the parametric modulator “prediction error” on the regressor “end of feedback presentation after the first trial” and checked which voxels had significant beta-values using FDR correction for multiple comparisons. Crucially, we also tested whether there was a significant interaction effect between time (first vs. second half of the task) and environment (low vs. medium vs. high noise environment) on the parametric modulator “prediction error” on the regressor “end of feedback presentation after the first trial” in one of the seven ROIs described above (six OFC subregions and the ventral striatum) using repeated measures ANOVAs and, if applicable, follow-up paired one-tailed t-test to interpret significant interaction effects. We used the same approach to investigate the effect of prediction errors on brain activity during feedback presentation on the second trial of each block.

For analysing the reward localiser, we used the same approach as for analysing the main task up to and including first-level analyses, where we used the first three events of interest from the main task. This resulted in 13 whole brain maps of parameter estimates (betas; three events of interest * two locations around the island + seven nuisance regressors). Next, we simply took the univariate contrast between the high and the low reward conditions at the first event of interest (i.e., island presentation). We did this because this should have shown us which brain regions were responsive to expected reward during the event of interest that our main analyses were focused on. In that way, we could control for this effect there. However, no brain regions were significantly responsive to reward expectancy during our reward localiser.

